# Long-read sequencing reveals telomere inheritance patterns from human trios

**DOI:** 10.1101/2025.10.07.680721

**Authors:** Yuxin Zhou, David R. Lougheed, Warren A. Cheung, Isabelle Thiffault, Tomi Pastinen, Guillaume Bourque

**Affiliations:** McGill University, Human Genetics, Montréal, QC H3A 0C7, Canada; Canadian Center for Computational Genomics, McGill University, Montréal, QC H3A 2R7, Canada; Children’s Mercy Hospital and Research Institute, Genomic Medicine Center, Kansas City, MO 64108, USA; Department of Pathology and Laboratory Medicine, Children’s Mercy Hospitals, Kansas City, MO 64108, USA; School of Medicine, University of Missouri-Kansas City, Kansas City, MO 64108, USA; Victor Phillip Dahdaleh Institute of Genomic Medicine at McGill University, Montréal, QC H3A 0G1, Canada; Institute for the Advanced Study of Human Biology (WPI-ASHBi), Kyoto University, Kyoto, 606-8303, Japan

## Abstract

Telomeres are essential for maintaining genomic integrity and are associated with cellular aging and disease, yet the factors influencing their inheritance across generations remain poorly understood. Leveraging PacBio HiFi long-read sequencing and 75 parent-offspring trios (n = 225) from the Genomic Answers for Kids program, we analyzed individual telomeres across chromosomes and their inheritance. Telomere length (TL) varied between chromosome arms in a way that was consistent in parents and offsprings, with average values ranging from 5000 to 8000 base pairs. Maternal and paternal TL together were a strong predictor of child TL (R^2^ = 0.59). Notably, using telomeric variant repeats, we developed a tool that enabled allelic tracing for 53.3% of maternally and 49.9% of paternally inherited telomeres. In the child, paternally transmitted alleles were significantly longer than age-matched maternal ones (Δmean = 409 bp, p = 2.6e-05), particularly when from older parents (Δmean = 698 bp, p = 8.9e-05) and at chromosome arms with shorter average TL (Δmean = 752 bp, p = 1.6e-06). These findings reveal parent-of-origin effects and heritable influences on TL, providing novel insights into telomere dynamics and their potential implications in age-related disease susceptibility.

## Introduction

Telomeres, composed of tandem TTAGGG repeats at chromosome ends, are essential for maintaining genome stability^1–3^. It is known that telomere length (TL) shortens with each cell division and can serve as a molecular indicator of replicative aging^4,5^. Critically short telomeres are implicated in rare disorders such as dyskeratosis congenita, where telomere dysfunction disrupts tissue renewal and hematopoiesis^6,7^. In contrast, long telomeres are observed in stem cells, immortalized cell lines, and tumors, which bypass replicative senescence through telomerase reactivation or alternative lengthening of telomeres mechanisms^8–10^.

In human, TL is influenced by both age-related decline and parental inheritance^11,12^. High heritability estimated for TL was reported in twin- and family-based studies^13,14^, and recent trio-based analysis confirmed that parental mean TL (MTL) can predict MTL in offsprings^15^. Paternal and maternal ages at conception were also found to have a significant contribution to offspring MTL^15–17^. In addition, evidence from mouse models has provided mechanistic insight into telomere inheritance. Indeed, parental MTL can determine offspring MTL even in the absence of telomerase, suggesting the existence of a fixed and heritable set point of telomere^18^. TL can also be actively remodeled in early embryos, with paternal TL influencing offspring TL^19,20^. Furthermore, recent reciprocal-cross experiments revealed a parent-of-origin effect in which the direction of embryonic TL change depends on the relative lengths of maternal and paternal telomeres^21^, indicating that telomere inheritance may be asymmetrically regulated depending on whether the longer or shorter telomere is inherited.

Although TL has been studied as a heritable trait, its inheritance dynamics at the level of individual chromosome arms and alleles remain largely unknown, especially in human. It is also unclear whether individual telomeres elongate after inheritance in a size-dependent manner. Addressing such questions could help reveal uncharacterized regulatory mechanisms but require more precise measurements of TL at the resolution of individual telomeres, which is not achievable with widely-used assays such as Flow-FISH, qPCR, TRF, and Southern blot^23–25^. Furthermore, even short-read sequencing approaches are challenged by the repetitive nature of telomeric and subtelomeric regions^27^. As a result, TL measurement at the resolution of individual chromosome arms and alleles has historically been technically constrained.

In contrast, long-read sequencing technologies such as Pacific Biosciences HiFi genome sequencing (HiFi-GS) and Oxford Nanopore Technologies now allow direct measurement of full telomeric regions, resolving TL and repeat content at each chromosome arm. Using such an approach, an enrichment-based long-read telomere profiling study in 147 individuals has shown consistent rank-ordering of chromosome arm-specific TLs established early in life and maintained with age^28^. The dynamic telomere shortening during replicative aging was also confirmed using long reads^29^. In addition to (TTAGGG)n repeats, long reads can also resolve the telomere variant repeat sequences (TVRs) in subtelomeric regions, which vary between homologous chromosome arms and individuals^30–32^. Telogator2, a tool to analyze telomeres from long-read data, measures allele-specific TL by clustering reads based on TVR composition^32^, and has been applied in a single-case family analysis to demonstrate TVR sharing between related individuals. However, long-read studies to date have not systematically examined telomere inheritance and whether it varies in a chromosome-arm-specific or parent-of-origin manner.

Here, we analyzed a cohort of 75 parent-child trios from the Genomic Answers for Kids (GA4K) program^33^ using HiFi-GS data. In addition to characterizing inter-individual TL variation across chromosome arms and the age-associated decline in TL, we analyzed family-based effect on TL, including the heritability of TL within trios using multivariate linear regression. To confirm the accuracy and clinical validity of our measurements, we benchmarked long-read TL estimates against Flow-FISH values in three pediatric probands with suspected telomere biology disorders. We then leveraged TVRs to compare alleles between parents and children, and developed a tool, named TeloScore, to identify directly inherited telomeric alleles. Using TeloScore, we tested for telomere drive, defined as the preferential transmission of longer or shorter telomeric alleles, and further examined whether the length and parental origin of inherited alleles contribute to TL variability in the child.

## Results

### 1. Telomere length varies significantly by chromosome arm

To characterize the variation of TL in the GA4K cohort, we analyzed the telomeres of 225 individuals from the 75 parent-offspring trios (Methods). Telomeric alleles were identified using Telogator2^32^, which detects canonical (TTAGGG)n repeats and TVRs in adjacent subtelomeric regions (Fig. 1a). Across the cohort, most chromosome arms had more than 800 telomeric reads (Supplementary Fig. 1a), which were usually resolved by Telogator2 into one or two TVR allele clusters (Supplementary Fig. 1b). TL was estimated only for allele clusters with sufficient telomeric read support (>3 reads), and arms without estimates were omitted. The allelic TL distribution for a single participant is shown in Fig. 1b. Overall, TL showed normal distribution patterns across individuals, with peak frequencies ranging from approximately 5000 to 8000 base pairs (Supplementary Fig. 1c). We divided chromosomal arms into three groups based on ranked median TL, with median TLs ranging from approximately 4800 to 5600 bp in short arms, 5600 to 6800 bp in medium arms, and 6900 to 8000 bp in long arms (Fig. 1c). Overall, the chromosome arm order based on median TL was found to be well preserved across children, mothers, and fathers (Supplementary Fig. 1d), with strong pairwise correlations observed between roles (Supplementary Fig. 1e). Notably, the median TL ordering also showed overall concordance to the results reported by Karimian et al.^28^ (Fig. 1d), even though those TLs were estimated using a different long-read enrichment-based approach.

**Fig 1.**
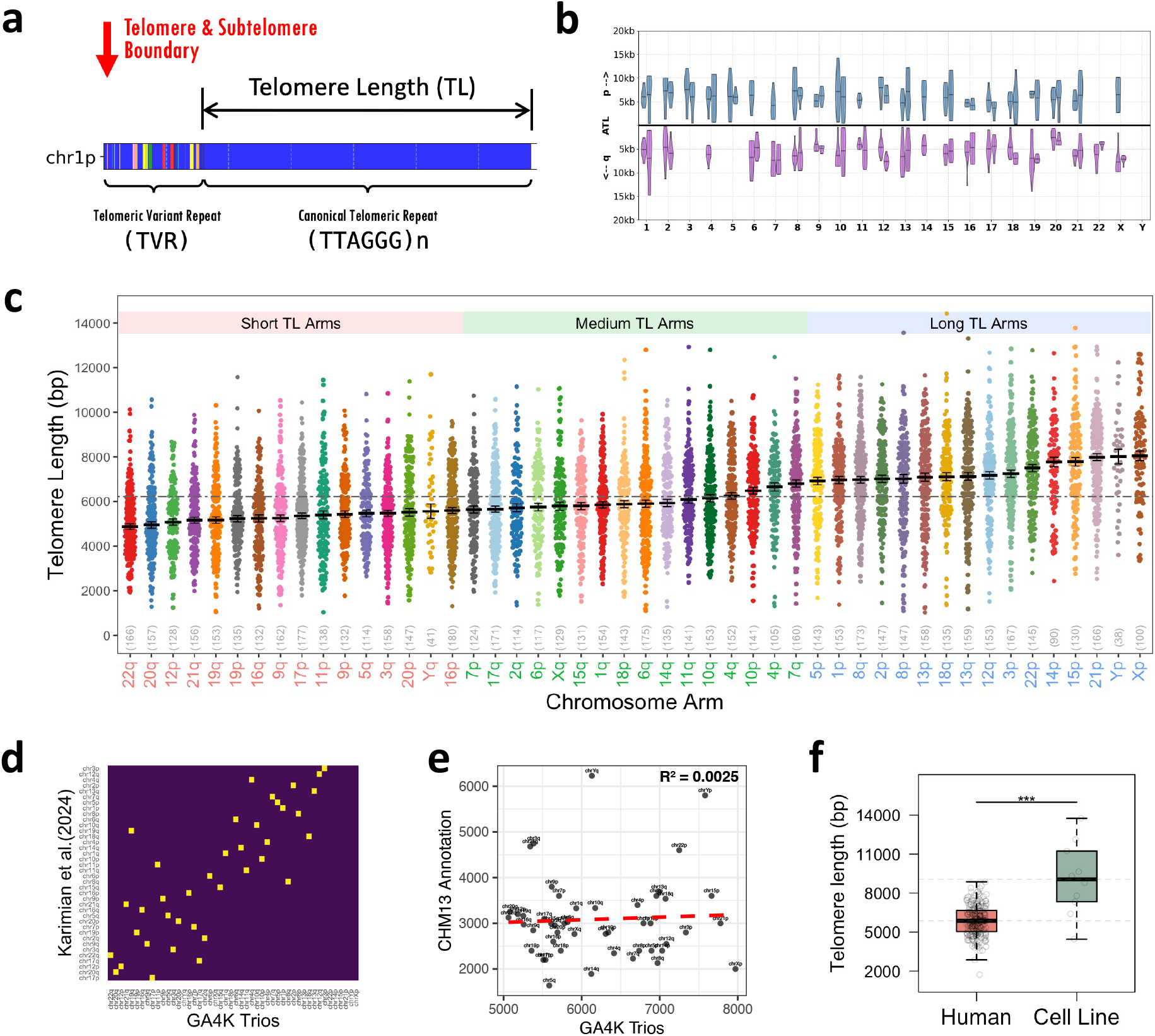
Telomere length varies significantly by chromosome arm. a. Telomere region composition. Telomeric variant repeat (TVR) and canonical telomeric region are shown chromosome 1p from one individual. b. Allelic TL of one individual (ID:cmh003312-01), directly obtained from the Telogator2 tool. c. Chromosome arm-specific TLs in the GA4K trio cohort (n = 225). Violin plot shows TLs for all 48 Chromosome arms, arranged in ascending order of median TL. Each data point corresponds to one individual’s estimated TL for a specific arm. Chromosomal arms are color-coded into short (red), medium green), and long (blue) groups. Black dashed lines indicate the median TL per arm, and whiskers represent the standard error of the mean. The gray dashed line indicates the overall mean of the median TLs across all arms. d. Arm-specific TL order comparison between GA4K trios and data reported by Karimian et al. (2024) in Science. The x-axis shows the chromosome arms in ascending order of median TL from left to right, as measured in the GA4K trio cohort; the y-axis shows the ascending order of median TL from bottom to top, as reported by Karimian et al. e. Scatter plot comparing the median arm-specific TL measurements in the GA4K cohort (n = 225) with those from the CHM13 reference genome annotation. Each point represents the median TL for a single chromosome arm in GA4K, plotted against the corresponding value in CHM13. f. Comparison of the MTL in GA4K trios versus Human Pangenome Reference Consortium (HPRC) cell lines.

Finally, we compared the GA4K trios with the telomeric annotation of the CHM13 T2T reference^34^, which showed little correlation (Fig. 1e). Average TLs was also found to be significantly longer in ten HiFi-sequenced lymphoblastoid cell lines from HPRC (Human Pangenome Reference Consortium)^35^ (Fig. 1f). In sum, we find that TL varied consistently between chromosome arms from both parents and offsprings, and that the arm-specific TL patterns observed in these blood samples were different from the ones observed in the assembled T2T reference or immortalized cell lines.

### 2. Telomere length is influenced by both age and parental inheritance

Having access to trio data, we wanted to explore the parental and age factors that contribute to offspring TL. For this analysis, we focused on MTL across all chromosome arms, since that is what traditional approaches have focused on^25^. We first compared MTLs across sample groups and family roles for the 68 parent-offspring trios with complete age and sex information (Fig. 2a-b). Offspring consistently showed longer MTLs than parents (p=3e-14) with no significant differences between males and females (p>0.05). Across all individuals, MTL declined markedly with age, with a strong negative correlation (R^2^=0.3268) (Fig. 2c). We also tested whether HiFi-GS coverage influenced to variation in TL estimates and observed only modest correlations (R^2^ = 0.033 for children and 0.083 for parents; Supplementary Fig. 2a). These results suggest that biological variation, including age, contributes more to TL differences across individuals than technical variation from sequencing coverage.

**Fig 2.**
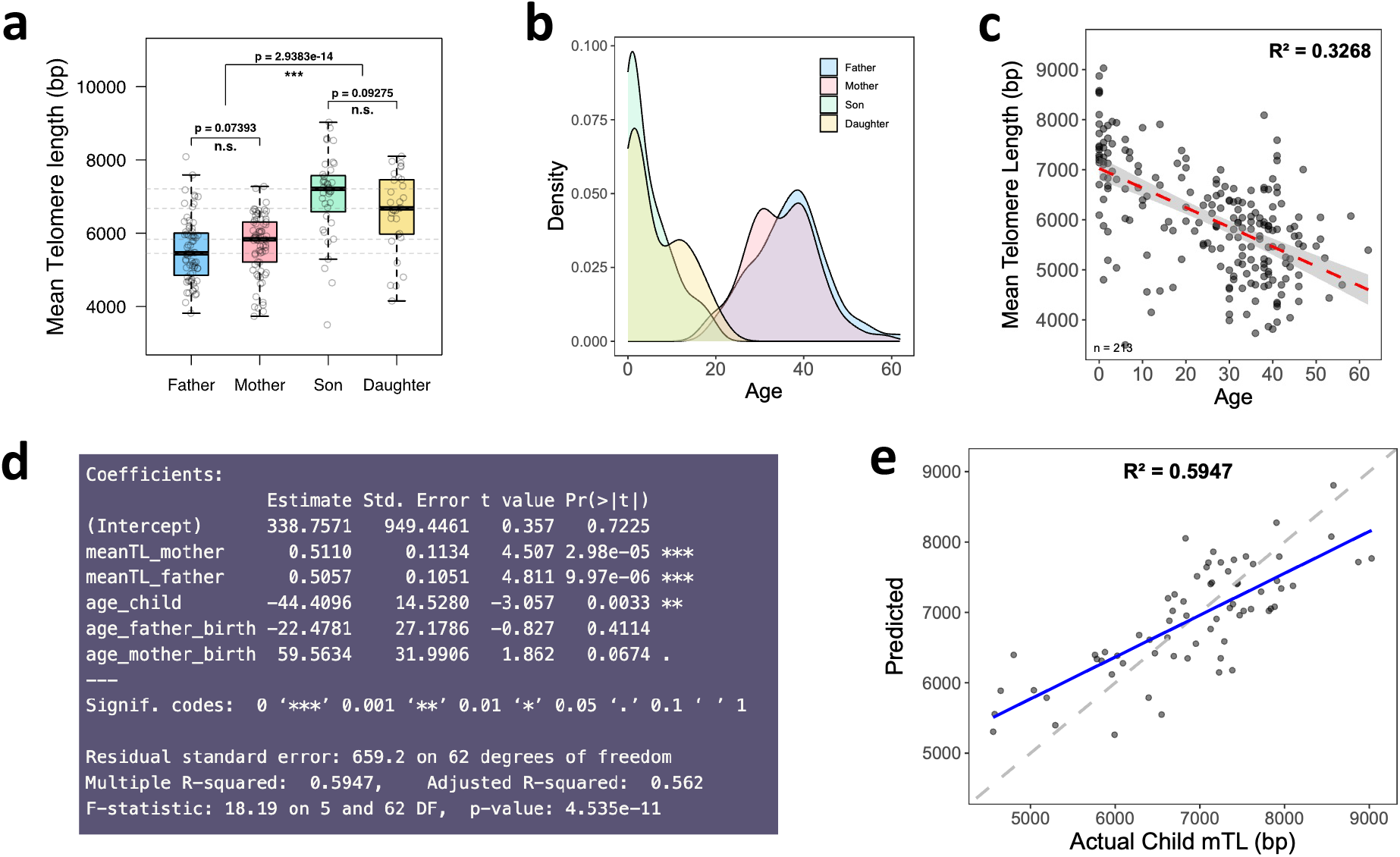
Telomere length is influenced by both age and parental inheritance. a. Comparison of mean telomere length (MTL) across family roles and sexes. Boxplots show variations in MTL among fathers, mothers, sons, and daughters, with statistical significance of differences evaluated between (1) fathers and mothers, (2) sons and daughters, and (3) parents and offspring, using the Wilcoxon rank-sum test (71 trios, n = 213). b. Age distribution across family roles in the GA4K cohort (71 trios, n = 213). c. Relationship between age and MTL for all participants (71 trios, n = 213), with Pearson’s R indicating a negative correlation. d. Summary of multivariate linear regression analysis showing significant predictors of offspring MTL. e. Observed offspring MTL and values predicted by the final model (69 trios, n = 204).

Next, to quantify the relative influence of parental MTLs and age-related factors on offspring TL, we applied a multivariate regression model, incorporating five predictors: parental MTLs, the child’s age, and the parental ages at conception (Methods). In this five-variable model, we found both maternal and paternal MTL were significantly contributing to child’s MTL (p-values of 2.98e-05 and 9.97e-05, respectively), and more so than maternal and paternal age (p-values of 0.0674 and 0.4114, respectively) (Fig. 2d). The model showed a fit achieving an R^2^ of 0.5947 (p=4.535e-11), surpassing the multiple R^2^ of 0.45 reported previously in an analysis of 250 trios using short-read sequencing^15^, which suggests an improved predictive performance with long-read data. Comparison of predicted and observed child MTLs showed close agreement overall, although the model tended to underpredict children with short telomeres (Fig. 2e). We also included sequencing coverage as a predictor to check for technical effects, showing that did not improve model fit (Supplementary Fig. 2b).

Having access to arm-specific TL measurements, we sought to test whether different sets of chromosome arms contributed differently to the predictive performance. We compared models using MTLs calculated across all arms, autosomes only, or excluding acrocentric arms to evaluate the impact model performance (Supplementary Table 1). Across all tested regression models, both maternal and paternal MTLs remained significant predictors of child MTL, with R^2^ values around 0.5 and overall model p-values < 1e-7. In most cases, maternal age at conception showed a stronger association than paternal age. Next, we tested whether the parental and age effects observed at the MTL level persist within each chromosome arms length category (short, medium and long, Fig. 1c). We found that age-related TL decline was evident in all groups (Supplementary Fig. 3a), with short-TL arms showing the most pronounced correlations, significantly exceeding those in the long-TL group (p=0.0075). Applying the five-variable model to each category revealed distinct parental contribution patterns (Supplementary Fig. 3b): short-TL arms were more strongly associated with maternal TL, medium arms showed a stronger paternal effect, and long arms displayed significant contributions from both parents, with the paternal effect being largest. Model fits were similar across categories, with R^2^ = 0.3908 in arms with short TLs, 0.3341 in arms with medium TLs, and 0.3566 in arms with long TLs.

**Fig 3.**
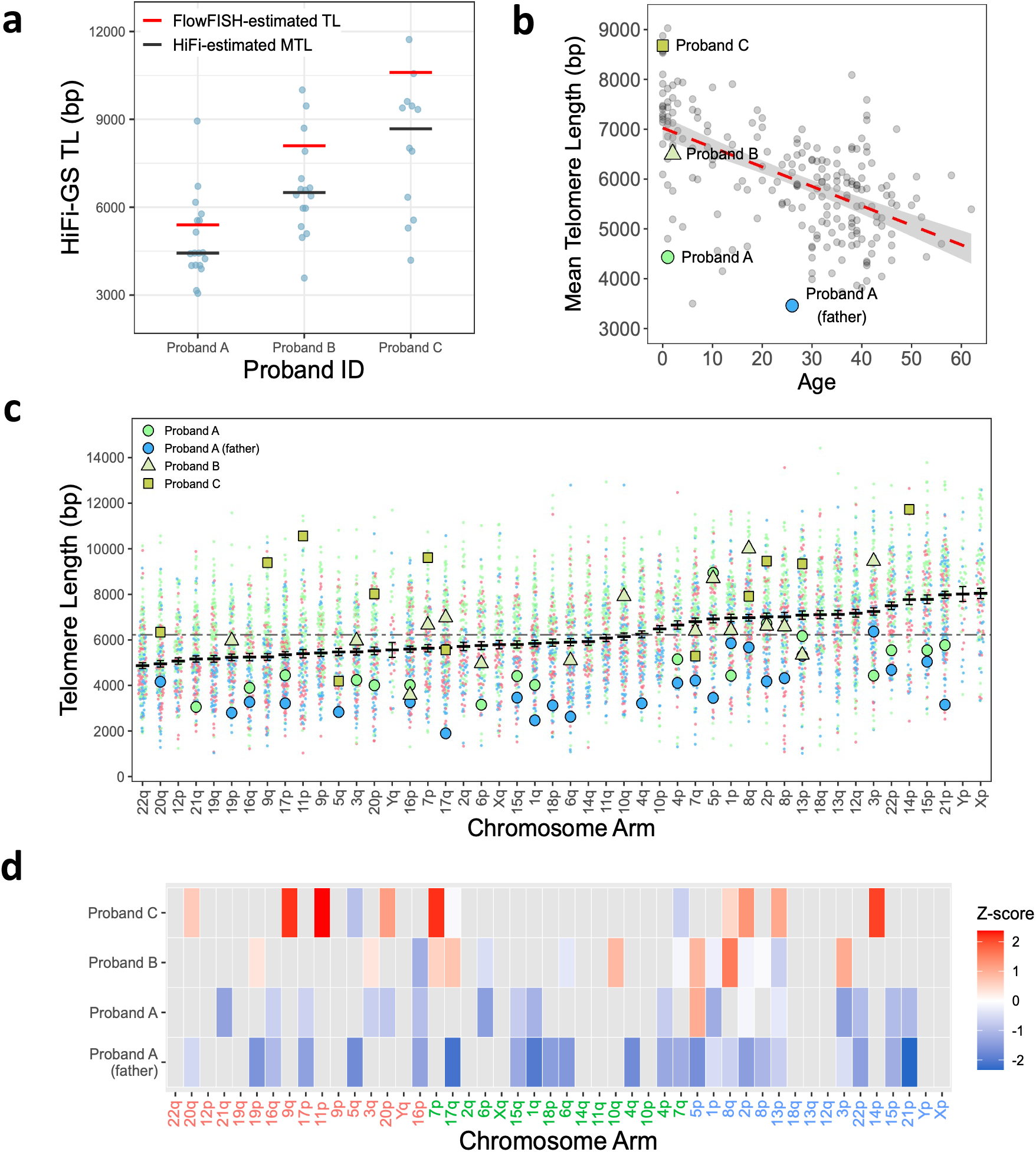
Arm-level telomeric shortening in pediatric patients with suspected telomere biology disorders. a. Chromosome-arm-specific TL estimates from HiFi-GS for each proband. Each blue point represents one arm TL measurement, with the black horizontal line showing the mean TL from the long reads. Red horizontal lines indicate TL measurements reported by Flow-FISH from lymphocytes. b. Comparison of HiFi-GS TLs in three probands and the father of Proband A against the age-matched distribution in the GA4K cohort (n = 204). c. Chromosome-arm-specific TLs for Proband A and his father, Proband B, and Proband C, compared to all children (green) and all fathers (blue) in the GA4K cohort (n = 225). Arms are ordered from shortest to longest based on cohort median TL. Both child and father show consistently reduced TL across chromosome arms relative to role-matched population TL distributions. d. Chromosome-arm-level TL Z-scores for three pediatric probands. Heatmap shows arm-specific TL Z-cores relative to the age-matched population distribution. Gray indicate chromosome arms with missing TL estimates. E. Arms are ordered from shortest to longest based on cohort median TL. Proband A displays widespread telomere shortening, whereas Proband B and Proband C show more variable patterns across chromosome arms.

Overall, we find that offspring MTL is robustly predicted by age and both parents’ MTL, and this relationship is independent of sequencing coverage. The best model fit was achieved when averaging across all chromosome arms, and model performance remained comparable across short, medium, and long arm categories, although the relative contributions of maternal and paternal TL differed between chromosome-arm groups.

### 3. Arm-level telomeric shortening in pediatric patients with suspected telomere biology disorders

To further validate the accuracy of our TL measurements and explore the clinical relevance of long-read-based telomere analysis, we analyzed three pediatric probands with suspected telomere biology disorders. All three probands underwent both PacBio HiFi-GS and clinical TL testing by Flow-FISH (Methods), enabling direct comparison between the two methods. Proband A additionally had a matched paternal sample, allowing evaluation of potential familial TL patterns. Clinical findings, genetic variants, and TL assessments for each proband are summarized in Supplementary Table 2. We compared MTL estimates obtained from long-read sequencing and Flow-FISH (Fig. 3a) and found that the two methods were consistent at the mean level, while long-read data additionally revealed the distribution of TL across chromosome arms. Using the age-MTL relationship from the GA4K cohort as a reference, we found that Probands B and C had MTLs within the expected age-matched range, whereas Proband A ranked among the lowest values for their age, a pattern also observed in the father of Proband A (Fig. 3b).

Next, we asked whether patterns of telomere shortening in the probands were localized to specific chromosome arms, which is a level of resolution not captured by traditional measurements. Comparing these proband’s arm-specific TLs with GA4K distributions from unaffected children and fathers revealed that Proband A and his father had consistent arm-level shortening, with values below the role-matched medians for most chromosome ends (Fig. 3c). Such distributed TL reductions were not observed in Probands B and C.

To further identify arms preferentially affected by telomere shortening, we calculated each arm’s Z-scores by comparing to role-matched samples without suspected telomere biology disorders (Fig. 3d). Proband A showed lower TL across multiple chromosome arms, with the strongest effects observed on 6p, 3p, and 21q. Although Proband B and Proband C appeared normal based on the mean TL, arm-specific measurement indicated TL reduction on 16p and 13p and 5q and 7q, respectively. The father of Proband A also exhibited shortened TL relative to his age and showed the strongest reductions on 21q, 17p, and 1q. The affected arms were distributed across all TL groups (short, medium, and long), suggesting that shortening was not limited to a specific group. Among all, chromosome 16p appeared consistently shortened in Proband A, his father and Proband B, yet measurements for Proband C were incomplete at this arm. While few arms showed overlapping shortening between Proband A and his father, both exhibited distributed arm-level TL reductions.

In sum, we find that long-read sequencing detects arm-specific TL reductions in pediatric probands with suspected telomere biology disorders, with Proband A and his father showing shortening across chromosome arms spanning both long and short TL groups, highlighting the clinical value of arm-level analysis for identifying telomere abnormalities that conventional methods do not detect.

### 4. TVR similarity identifies inherited telomeric alleles

Having observed TL associations between parents and offspring, we aimed to determine how individual telomeric alleles are inherited and what factors contribute to this process. We developed a custom scoring framework to quantify TVR similarity between alleles, which we called TeloScore (Methods), to allow allele-level tracking between parents and children. With TeloScore, a similarity score between 0 and 1 was computed for each pair of TVR sequences, with higher values indicating closer structural matches of variant repeat patterns. Before applying this method to parent-child comparisons, we first evaluated TVR similarity patterns across unrelated individuals to assess how much variation exists at each chromosome arm. Most arms showed high variability between individuals, whereas 2p was an outlier with consistently high similarity (Supplementary Fig. 4a,b). Analyses of the number of distinct TVR allele clusters and the distribution of TVR region lengths indicated that these features did not explain the 2p pattern (Supplementary Fig. 1b and Supplementary Fig. 4c,d). Notably, in parent-child trios, the TeloScore allowed to assign child alleles to their parental origin. In a representative family, child alleles on chromosome arm 13q showed one-to-one matches to both maternal and paternal alleles (Fig. 4a). In this case, both the best matched maternal allele and the best paternal allele scored 1.0 (Fig. 4b). Across all reported alleles in this trio, the full similarity matrix exhibited a strong diagonal (Fig. 4c), showing consistent direct inheritance on a per-arm basis.

**Fig 4.**
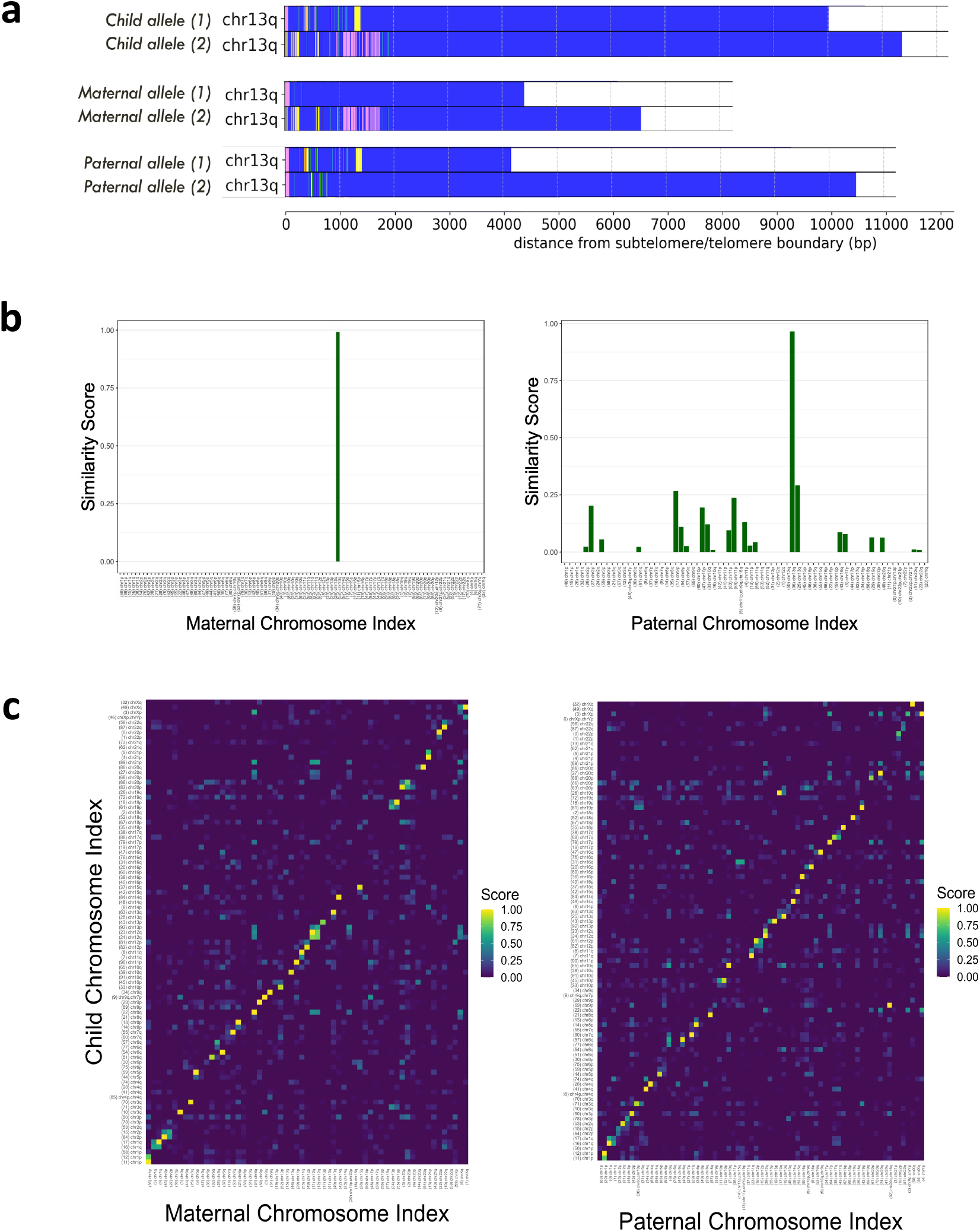
TVR similarity identifies inherited telomeric alleles. a. Visualization of TVR structures for all six alleles (two child, two maternal, two paternal) at the 13q chromosome arm in a representative trio. Repeat structures are aligned from the subtelomere-telomere boundary (position 0), with colored bars representing different telomeric k-mers and their spans. One child allele shows a clear structural match to a parent, supporting direct inheritance. b. Similarity scores comparing one child allele to all maternal (left) or paternal (right) alleles across arms. In most cases, only one parental allele reaches high similarity (score close to 1.0), enabling confident assignment of the child allele to its parental origin. c. TVR similarity heatmaps comparing all child alleles to maternal (left) and paternal (right) alleles in a representative trio. A strong diagonal indicates consistent one-to-one inheritance, where each child allele is most similar to one specific parental allele at the same chromosome arm.

Overall, we had telomeric alleles on 3026 arms in children, 1909 in mothers, and 1961 in fathers across all trios (Supplementary Table 3). To define a cutoff for calling a TVR match as inherited using TeloScore, we examined the distribution of TVR similarity scores across all parent-child chromosome arm comparisons. Scores showed a strong peak near 1.0, and most comparisons between different chromosomes scored much lower (Supplementary Fig. 4e). A similarity threshold of 0.7 was selected to call inherited TVR patterns with high confidence. At this cutoff, the error rate was estimated to be below 3% based on the number of predicted inherited alleles with an incorrect arm assignment (Supplementary Table 4). With this approach, among 1593 chromosome arms where both mother and child had telomeric allele calls, we identified 849 cases (53.3%) where the child’s allele matched one from the mother. Similarly, among 1618 arms shared between father and child, 807 inheritance matches (49.9%) were successfully identified (Supplementary Table 4). Overall, TeloScore allowed confident identification of inherited telomeric alleles using variant repeat sequences, enabling investigation of factors contributing to inheritance at the allelic level.

### 5. Direct inherited TLs reveal transmission asymmetries and parent-specific effects

Finally, we examined telomere inheritance at the chromosome arm and allele levels by assessing transmission preference and parent of origin’s influences. We first tested for telomere drive, defined as a bias in the transmission of one of the two parental alleles at a given chromosome arm to the offspring. In total, 612 parent-child chromosome-arm pairs (313 mother-child and 299 father-child) had two alleles TL measurements in the parent (Supplementary Table 3) and a predicted inherited allele in the child. We then calculated the difference between the transmitted allele’s TL, and the average TL of parental alleles present at that arm (Methods). Allele-specific telomere transmission bias was centered around zero (Fig. 5a), suggesting randomized transmission and the absence of telomere drive. This pattern remains consistent after stratifying by parental origin, showing no transmission bias from either mother or father to offspring (Supplementary Fig. 5a).

**Fig 5.**
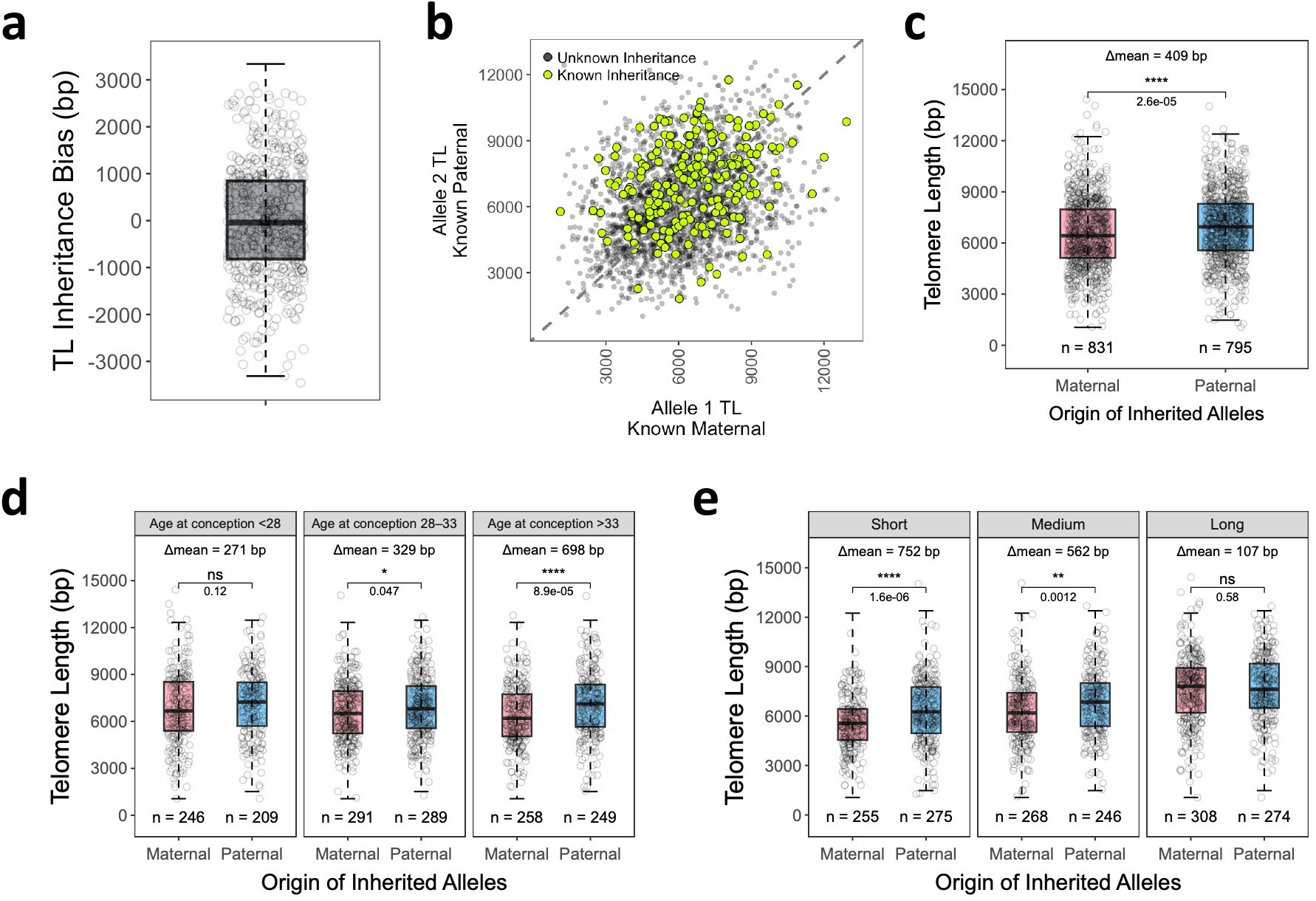
Direct inherited TLs reveal transmission asymmetries and parent-specific effects. a. Telomere drive analysis of transmitted alleles. Inheritance bias was defined as the difference between he inherited allele TL and the average TL of both alleles from the transmitting parent at the same chromosome arm. No consistent bias was observed, indicating no preferential transmission of longer or shorter alleles. b. Lengths of telomeric allele pairs detected at the same chromosome arm in each child. Gray data points represent two alleles detected at a given chromosome arm without assigned parental origin, whereas green data points represent child chromosome arms where maternal and paternal origin were determined using TeloScore. c. Comparison of inherited telomeric alleles transmitted from paternal and maternal origin. Paternally transmitted alleles showed significantly longer TL than maternally transmitted alleles. Maternal- and paternal-origin alleles were analyzed as independent measurements in statistical tests. d. Stratification of the parent-of-origin difference in (C) by parental age at conception (<28, 28–33, >33 years). The TL difference between paternally and maternally transmitted alleles increased with parental age, with the largest difference observed in the oldest group. e. Stratification of the parent-of-origin difference in (C) by parental TL rank at each chromosome arm, using the short, medium, and long categories defined in (B). Paternally transmitted alleles had longer TL than maternally transmitted alleles across all categories, with the largest length difference and strongest significance observed in short arms, while the difference was not significant in long arms.

Inheritance is decided in the germ line, while our TL measurements are obtained from blood cells. To partially circumvent this caveat, we next focused on the child telomeres, directly comparing maternally and paternally inherited alleles to assess whether inherited TL differs by parent of origin. Figure 5b shows the distribution of TL between allele pairs detected at the same chromosome arm in each child, overlaid with TL values of alleles with known parental origin. To test whether a difference could be observed, we grouped all directly inherited alleles in the child by parental origin and compared their TL distributions. Notably, paternal-origin alleles were significantly longer than maternal-origin alleles in the child (Δmean =409 bp, p = 2.6e-05) (Fig. 5c). This pattern was also consistent across chromosome arms (Supplementary Fig. 5b), with paternal-origin alleles remaining longer telomere than maternal-origin alleles in the child. This result contrasts with the paternal telomeres, measured in the blood, which are typically shorter than their maternal counterparts (Fig. 2a).

To further explore telomere inheritance in the child, we tested whether parental age at conception modulates the TL difference between maternal-origin and paternal-origin alleles. We grouped inherited telomeric alleles by parental age at conception (i.e., age of the transmitting parent), and compared the TL of paternal- and maternal-origin alleles within each group. Each allele was treated as an independent data point. Children’s telomere alleles showed similar TL in the youngest parental age-at-conception group (<28), but paternal-origin alleles became progressively longer than maternal-origin alleles in older groups (28–33 and >33 years), reaching an average 698 bp difference in trios with parental age at conception above 33 years (Fig. 5d). Therefore, telomeric alleles of paternal origin from older fathers at conception are maintained at longer lengths compared to those of maternal origin.

Lastly, to assess whether parent-of-origin effects on telomere transmission vary by chromosome arms, we grouped inherited alleles according to the short, medium, and long arm categories defined based on the full cohort (Fig. 1c). Paternally inherited alleles were longer than maternally inherited alleles in all TL arm groups, with significant differences in the short (Δmean = 752 bp, p = 1.6e-06) and medium (Δmean = 562 bp, p = 0.0012) groups, but not in the long group (Δmean = 107 bp, p = 0.58) (Fig. 5e). The largest difference occurred in short arms, where paternal-origin alleles exceeded maternal-origin alleles by approximately 1200 bp on average. This suggests that parent-of-origin effects are most pronounced in chromosome arms with shorter TLs. In sum, parent-specific effects on telomere inheritance are influenced by both parental age at childbirth and chromosome-arm-specific TL.

## Discussion

Long-read HiFi-GS from 225 whole blood samples revealed that human telomere varies substantially between chromosomal arms. This arm-specific pattern aligns with recent findings using enrichment-based long-read sequencing from human PBMCs^28^. Within the trios we profiled, the TL rank ordering was preserved across different family roles (i.e., mother, father, and child), implying a stable regulation of TL at the chromosome arm level. However, such inter-arm TL pattern was not reflected in current reference assemblies including Telomere-to-Telomere (T2T) and Human Pangenome, which are typically derived from immortalized cell lines with distorted telomere profiles. This highlights the need to profile appropriate cell types to study telomere variation and patterns of inheritance.

We confirmed that parental MTLs and child age together provide a meaningful prediction of the offspring’s MTL, highlighting both the contribution from each parent and the effect of age-related telomere shortening. Although our model achieved higher predictive accuracy than previous trio-based study using NGS^15^, we observed that the model tended to underpredict TL in children with particularly short telomeres. When we stratified the model by TL group (short, medium, or long arms), predictive performance decreased, likely due to reduced sample size within each group. Expanding the trio dataset could improve modeling accuracy, especially for individuals with particularly short or long TL, yet model performance might also be limited by the fact that we are using blood samples to approximate germline inheritance.

Pathology in telomere biology disorders could arise from a single critically short telomere, highlighting the clinical relevance of high-resolution arm-specific TL measurement^36^. Long-read sequencing enabled arm-level TL analysis in patients with suspected telomere biology disorders, revealing patterns not captured by MTL or standard Flow-FISH assays. In particular, Proband A and his father both exhibited distributed shortening across specific chromosome arms, and some affected arms were recurrently observed across multiple families. These findings suggest that arm-specific TL vulnerability may underlie certain clinical presentations of telomere biology disorders. Further studies with larger patient cohorts are needed to determine whether recurrent arm-level patterns exist and reflect distinct biological mechanisms or clinical subtypes.

By developing a telomere similarity scoring method (TeloScore), we were able to assign child telomeric alleles to the parent of origin and trace inheritance patterns in 50% of chromosome arms. We note that the number of matched alleles was limited, partly because Telogator2 clustering step occasionally produced more than two candidate alleles, making it difficult to determine the correct match. Refining allele clustering and TVR region definitions should increase the number of informative arms and improve inheritance tracking.

Nonetheless, using the 1656 parent-child inherited alleles we could assign with confidence, we observed a persistent parent-specific asymmetry in direct telomere inheritance. In the child, paternally inherited alleles were consistently longer than maternally inherited ones, and this asymmetry was more pronounced in children of fathers with older age at conception. These patterns are not explained by telomere drive of longer or shorter alleles but instead suggest that both germline TL and early embryonic processes contribute to determining telomere inheritance in the offspring. Unlike oogenesis, human spermatogenesis retains telomerase activity, and sperm TL increases with paternal age^37^, which could lead to the transmission of longer telomeres in older fathers. Notably, the observed maternal-paternal asymmetry was also stronger at chromosome arms with shorter median TLs. One potential explanation is that the elongation in the paternal germline preferentially targets shorter telomeres. Studies in yeast and human cells have shown that short telomeres are preferentially elongated by telomerase, while longer telomeres are actively suppressed by length-dependent feedback mechanisms^38^.

Together, these findings demonstrate that telomere inheritance is not simply a matter of parental TL averages. Instead, it reflects an interplay between the length of transmitted telomeric alleles and their parental origin, influenced both by germline and embryonic mechanisms at an allele-specific level. Additional studies will be needed to further shed light on these processes and perhaps uncover some of the regulatory mechanisms explaining the differences observed in chromosome arms that have typically short, medium or long telomeres.

## Methods

### Human trio cohort

Whole blood samples were collected from 225 individuals forming 75 parent-offspring trios enrolled in the Genomic Answers for Kids (GA4K) program at Children’s Mercy Hospital. Each trio consists of a child and both biological parents. DNA was extracted from whole blood using the chemagic™ 360 automated platform (PerkinElmer)^39^. Age information was available for 213 participants across 71 trios. Sex was recorded at enrollment and confirmed via genomic data (48% female, 52% male). Proband age was defined at the time of sequencing. Reported ages ranged from 0 to 21 years for children (median: 2), 19 to 58 years for mothers (median: 35), and 20 to 62 years for fathers (median: 37).

### Children with suspected telomere biology disorders

Three probands with suspected telomere biology disorders were included in this study. Proband A had a matched paternal sample included for analysis. Genomic findings include a 10.6 Mb deletion at 14q13.2–14q21.2 and a heterozygous TERT mutation (c.3187G>A, p.G1063S) in Proband A; a de novo X-linked DKC1 variant (c.977G>T, p.Gly326Val) and abnormal X-inactivation in Proband B; and a de novo DOT1L mutation (c.367G>A, p.Glu123Lys) in Proband C, associated with autosomal dominant neurodevelopmental disorders.

### Cell line dataset

Public PacBio HiFi-GS data from 10 lymphoblastoid cell lines were obtained from the Human Pangenome Reference Consortium (HPRC). These datasets were sequenced using the Sequel II platform with circular consensus sequencing (CCS v4.0.0), and the average coverage was approximately 39×.

### HiFi genome sequencing

All GA4K samples, including the three probands with suspected telomere biology disorders, were sequenced using PacBio HiFi genome sequencing (HiFi-GS). DNA from each sample was sequenced using the Sequel II system with circular consensus sequencing (CCS). Sequencing depth averaged ∼25× for children and ∼10× for each parent. Multiple runs were merged per individual and processed using the PacBio WGS WDL pipeline with default parameters^35^.

### TL estimation from long reads

We used Telogator2^32^ to estimate TLs from HiFi-GS reads. Telomeric candidate reads were identified based on telomeric k-mer density. The sub-telomeric regions of these reads were aligned with Winnowmap^41^ to the Telogator2 custom reference genome, which contains several reference subtelomere regions of same length, using the command: *python telogator2*.*py -i hifi*.*fa -o results/ --winnowmap winnowmap*. For arm-level TL analysis, we extracted the portion of each read containing only canonical TTAGGG repeats and calculated the 90th percentile of lengths per arm. Mean telomere length (MTL) for each individual was computed by averaging canonical TLs across all reported chromosome arms. TVR sequences adjacent to the canonical telomere region were also extracted using Telogator2 with default settings and encoded as k-mer consensus strings with single-letter symbols.

### Clinical Flow-FISH TL assay

Probands diagnosed with suspected telomere biology disorders underwent RepeatDx® using a clinical Flow-FISH assay, which reports median TLs in lymphocytes and other leukocyte subsets. The assay is accredited in North America and based on a validated protocol^42^. To benchmark long-read TL estimation, we compared Telogator2-derived TLs from whole blood with corresponding Flow-FISH values. TLs based on canonical repeats showed a consistent offset of ∼1 kb relative to Flow-FISH, which was reduced when including adjacent variant repeats (TVRs).

### Multivariate regression modeling of child TL

We used R’s *lm()* function to fit a multiple linear regression model predicting child MTL from five predictors: *MTL_child ∼ MTL_mother + MTL_father + Age_child + Age_mother_at_conception + Age_father_at_conception*.Parental ages at childbirth were included based on prior trio studies using next-generation sequencing^15^ and published evidence on parental age effects^37^. Prior to regression modeling, we excluded four trios with unknown age. Additionally, two other trios were omitted as the child’s TL was unexpectedly short compared to parental values, and a separate trio was excluded due to a child TL more than twice the parental average. The final model included 204 individuals from 68 trios.

### Computing TVR similarity using TeloScore

To quantify the similarity between two TVR alleles, we developed TeloScore, an in-house comparison-scoring tool. The tool calculates pairwise similarity scores between TVR consensus strings and generates a similarity matrix where each entry represents the alignment score between a pair of TVR consensus alleles. Scores range from 0 (very little similarity) to 1 (perfect match) and are calculated using a custom version of the Needleman-Wunsch (NW) alignment scoring algorithm^43^ which prioritizes TVR-specific repeat patterns and discounts shared canonical repeats within the TVR sequence. To compute this score, TeloScore makes use of the NW implementation in the *parasail* library^44^ and applies the following high-level algorithm:

1. Compress runs of the same TVR repeat motifs in the TVR sequence, using the logarithm of the length instead, to reduce the weight of length variations in these runs.
2. Apply the NW algorithm using a custom scoring matrix which de-emphasizes matches between canonical telomere repeat motifs to prioritize TVR-specific repeat patterns.
3. Adjust the score to be between 0 and 1, where 1 indicates a perfect match and 0 indicates little similarity between the sequences.

For these analyses, we used TeloScore v0.4.0 with scoring algorithm “1” and default parameter values.

### Allele-level inheritance analysis

We performed downstream analyses on inherited alleles identified based on TVR similarity using TeloScore. Allele-level TL analyses were restricted to chromosome arms with high-confidence inheritance matches. A child allele was considered directly inherited if it matched either a maternal or paternal allele with a TVR similarity score ≥ 0.7. For most chromosome arms, a single parent allele matched a single child allele above the similarity threshold, and this one-to-one match was used directly in the analysis. If multiple clusters from the same arm passed the similarity threshold, we treated them as one inherited allele and merged their telomeric reads. In cases where matching pattern could not be clearly resolved, such as when both parents showed high similarity to the same child allele, we marked the case as a conflict and excluded it from further analysis. If a parent allele showed its highest similarity to a child allele on a different chromosome arm, the pair was classified as an off-arm match and counted toward the estimated error rate.

To identify whether the longer or shorter parental allele is preferentially transmitted to offspring, we calculated the inheritance bias score, defined as the difference between the TL of the transmitted allele and the average TL of parental alleles on the same chromosome arm. Using the average provided a consistent baseline to compare the transmitted allele, especially in cases where more than two parental clusters were detected. This score quantifies whether the inherited allele is longer or shorter relative to the parental average. A positive bias indicates preferential transmission of the longer parental allele, whereas a negative bias suggests transmission of the shorter one.

## Data availability

GA4K HiFi-GS data analyzed in this study are available under controlled access via dbGaP and hosted on the AnVIL platform. Trios’ data are available through dbGaP Genomic Answers for Kids accession phs002206.v5.p1.

## Code availability

The source code for TeloScore is available on GitHub (https://github.com/davidlougheed/teloscore) and archived in Zenodo (DOI: 10.5281/zenodo.14984161).

## Author contributions

Y.Z., T.P., and G.B. conceived and designed the study. Y.Z. performed data analysis, and figure and table preparation. W.A.C. and I.T. coordinated biosample collection, sequencing, and quality control. D.L. developed the TeloScore algorithm and wrote the associated code, with input from Y.Z. T.P. and G.B. supervised the project. Y.Z., D.L., T.P., and G.B. contributed to data interpretation. Y.Z. and G.B. wrote the manuscript with input from all authors. All authors reviewed and approved the final manuscript.

## Acknowledgements

We thank all families for participating in the GA4K study. This work was made possible by generous gifts to the Children’s Mercy Research Institute and GA4K program at Children’s Mercy Kansas City. We also thank Nick Nolte, Dan Louiselle, and Rebecca Biswell for their work in sample processing, Laura Puckett and Adam Walters for their work in library preparation and sequencing, and the clinical coordination team led by Bradley Belden for their work in clinical coordination. Additional thanks to PacBio for sequencing support for a subset of the samples. T.P. holds the Dee Lyons/Missouri Endowed Chair in Pediatric Genomic Medicine, and B.G. holds the Roberta D. Harding & William F. Bradley, Jr. Endowed Chair in Genomic Research. G.B. is supported by a Canada Research Chair Tier 1 award and an FRQ-S Distinguished Research Scholar award. This research was enabled by a grant from the Natural Sciences and Engineering Research Council of Canada (RGPIN-2024-05329) and computational resources provided by Calcul Québec and the Digital Research Alliance of Canada. We are grateful to Dr. Cristian Groza for the technical support and all members of the Bourque lab and C3G group for their valuable discussion and supportive research environment. We also wish to thank Mia Levine and Michael Lampson, who helped us in the design of the study and provided us with feedback on the manuscript.

## Competing interests

The authors declare that they have no conflict of interest.

## Notes

### Competing Interest Statement

The authors have declared no competing interest.

## References

1. Greider, C. W. & Blackburn, E. H. Identification of a specific telomere terminal transferase activity in tetrahymena extracts. Cell 43, 405–413 (1985).

2. Moyzis, R. K. et al. A highly conserved repetitive DNA sequence, (TTAGGG)n, present at the telomeres of human chromosomes. Proceedings of the National Academy of Sciences 85, 6622–6626 (1988).

3. O’Sullivan, R. J. & Karlseder, J. Telomeres: protecting chromosomes against genome instability. Nat Rev Mol Cell Biol 11, 171–181 (2010).

4. Levy, M. Z., Allsopp, R. C., Futcher, A. B., Greider, C. W. & Harley, C. B. Telomere end-replication problem and cell aging. Journal of Molecular Biology 225, 951–960 (1992).

5. Allsopp, R. C. et al. Telomere Shortening Is Associated with Cell Division in Vitro and in Vivo. Experimental Cell Research 220, 194–200 (1995).

6. Mitchell, J. R., Wood, E. & Collins, K. A telomerase component is defective in the human disease dyskeratosis congenita. Nature 402, 551–555 (1999).

7. Kam, M. L. W., Nguyen, T. T. T. & Ngeow, J. Y. Y. Telomere biology disorders. npj Genom. Med. 6, 1–13 (2021).

8. Strahl, C. & and Blackburn, E. H. Effects of Reverse Transcriptase Inhibitors on Telomere Length and Telomerase Activity in Two Immortalized Human Cell Lines. Molecular and Cellular Biology 16, 53–65 (1996).

9. Blasco, M. A. Telomere length, stem cells and aging. Nat Chem Biol 3, 640–649 (2007).

10. Cesare, A. J. & Reddel, R. R. Alternative lengthening of telomeres: models, mechanisms and implications. Nat Rev Genet 11, 319–330 (2010).

11. Müezzinler, A., Zaineddin, A. K. & Brenner, H. A systematic review of leukocyte telomere length and age in adults. Ageing Research Reviews 12, 509–519 (2013).

12. Rossiello, F., Jurk, D., Passos, J. F. & d’Adda di Fagagna, F. Telomere dysfunction in ageing and age-related diseases. Nat Cell Biol 24, 135–147 (2022).

13. Slagboom, P. E. & Droog, S. Genetic Determination of Telomere Size in Humans: A Twin Study of Three Age Groups. Am. J. Hum. Genet. (1994) doi:1994%20 Nov;55(5):876–882.

14. Broer, L. et al. Meta-analysis of telomere length in 19,713 subjects reveals high heritability, stronger maternal inheritance and a paternal age effect. Eur J Hum Genet 21, 1163–1168 (2013).

15. Nersisyan, L., Nikoghosyan, M. & Arakelyan, A. WGS-based telomere length analysis in Dutch family trios implicates stronger maternal inheritance and a role for RRM1 gene. Sci Rep 9, 18758 (2019).

16. Unryn, B. M., Cook, L. S. & Riabowol, K. T. Paternal age is positively linked to telomere length of children. Aging Cell 4, 97–101 (2005).

17. Eisenberg, D. T. A., Hayes, M. G. & Kuzawa, C. W. Delayed paternal age of reproduction in humans is associated with longer telomeres across two generations of descendants. Proceedings of the National Academy of Sciences 109, 10251–10256 (2012).

18. Chiang, Y. J. et al. Telomere length is inherited with resetting of the telomere set-point. Proceedings of the National Academy of Sciences 107, 10148–10153 (2010).

19. Liu, L. et al. Telomere lengthening early in development. Nat Cell Biol 9, 1436–1441 (2007).

20. Frutos, C. de et al. Spermatozoa telomeres determine telomere length in early embryos and offspring. (2016) doi:10.1530/REP-15-0375.

21. Jeon, H.-J., Levine, M. T. & Lampson, M. A. A parent-of-origin effect on embryonic telomere elongation determines telomere length inheritance. Preprint at 10.1101/2025.01.28.635226 (2025).

22. Wang, J., Chen, P.-J., Wang, G. J. & Keller, L. Chromosome Size Differences May Affect Meiosis and Genome Size. Science 329, 293–293 (2010).

23. Baerlocher, G. M., Vulto, I., de Jong, G. & Lansdorp, P. M. Flow cytometry and FISH to measure the average length of telomeres (flow FISH). Nat Protoc 1, 2365–2376 (2006).

24. Alder, J. K. et al. Diagnostic utility of telomere length testing in a hospital-based setting. Proceedings of the National Academy of Sciences 115, E2358–E2365 (2018).

25. Montpetit, A. J. et al. Telomere Length: A Review of Methods for Measurement. Nursing Research 63, 289 (2014).

26. Graakjaer, J. et al. The pattern of chromosome-specific variations in telomere length in humans is determined by inherited, telomere-near factors and is maintained throughout life. Mechanisms of Ageing and Development 124, 629–640 (2003).

27. Treangen, T. J. & Salzberg, S. L. Repetitive DNA and next-generation sequencing: computational challenges and solutions. Nat Rev Genet 13, 36–46 (2012).

28. Karimian, K. et al. Human telomere length is chromosome end–specific and conserved across individuals. Science 384, 533–539 (2024).

29. Schmidt, T. T. et al. High resolution long-read telomere sequencing reveals dynamic mechanisms in aging and cancer. Nat Commun 15, 5149 (2024).

30. Tham, C.-Y. et al. High-throughput telomere length measurement at nucleotide resolution using the PacBio high fidelity sequencing platform. Nat Commun 14, 281 (2023).

31. Grigorev, K. et al. Haplotype diversity and sequence heterogeneity of human telomeres. Genome Res. (2021) doi:10.1101/gr.274639.120.

32. Stephens, Z. & Kocher, J.-P. Characterization of telomere variant repeats using long reads enables allele-specific telomere length estimation. BMC Bioinformatics 25, 194 (2024).

33. Cohen, A. S. A. et al. Genomic answers for children: Dynamic analyses of >1000 pediatric rare disease genomes. Genetics in Medicine 24, 1336–1348 (2022).

34. Nurk, S. et al. The complete sequence of a human genome. Science 376, 44–53 (2022).

35. Liao, W.-W. et al. A draft human pangenome reference. Nature 617, 312–324 (2023).

36. Rossiello, F., Jurk, D., Passos, J. F. & d’Addadi Fagagna, F. Telomere dysfunction in ageing and age-related diseases. Nat Cell Biol 24, 135–147 (2022).

37. Eisenberg, D. T. A. & Kuzawa, C. W. The paternal age at conception effect on offspring telomere length: mechanistic, comparative and adaptive perspectives. Philosophical Transactions of the Royal Society B: Biological Sciences 373, 20160442 (2018).

38. Britt-Compton, B., Capper, R., Rowson, J. & Baird, D. M. Short telomeres are preferentially elongated by telomerase in human cells. FEBS Letters 583, 3076–3080 (2009).

39. Cheung, W. A. et al. Direct haplotype-resolved 5-base HiFi sequencing for genome-wide profiling of hypermethylation outliers in a rare disease cohort. Nat Commun 14, 3090 (2023).

40. Farrow, E. et al. O24: Unveiling the power of HiFi genome sequencing: One test to rule them all? Genetics in Medicine Open 2, 101471 (2024).

41. Long-read mapping to repetitive reference sequences using Winnowmap2 | Nature Methods. https://www.nature.com/articles/s41592-022-01457-8.

42. Baerlocher, G. M., Vulto, I., de Jong, G. & Lansdorp, P. M. Flow cytometry and FISH to measure the average length of telomeres (flow FISH). Nat Protoc 1, 2365–2376 (2006).

43. Needleman, S. B. & Wunsch, C. D. A general method applicable to the search for similarities in the amino acid sequence of two proteins. Journal of Molecular Biology 48, 443–453 (1970).

44. Daily, J. Parasail: SIMD C library for global, semi-global, and local pairwise sequence alignments. BMC Bioinformatics 17, 81 (2016).

