## Supplemental Figures and Tables for "Long-read sequencing reveals telomere inheritance patterns from human trios"

Sfig.1. part 1/2

**A**

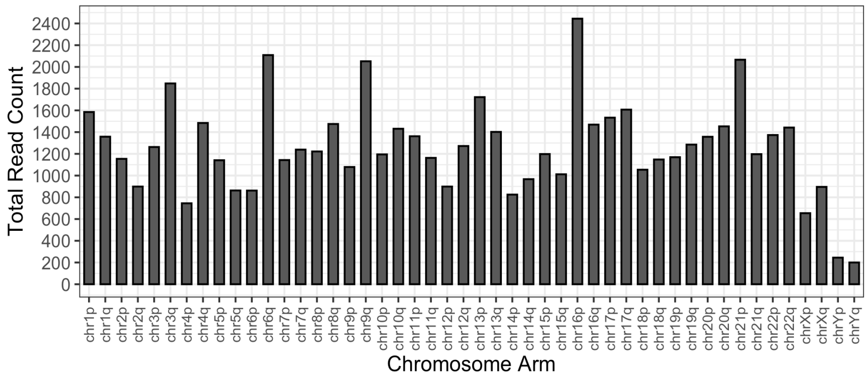


**B**

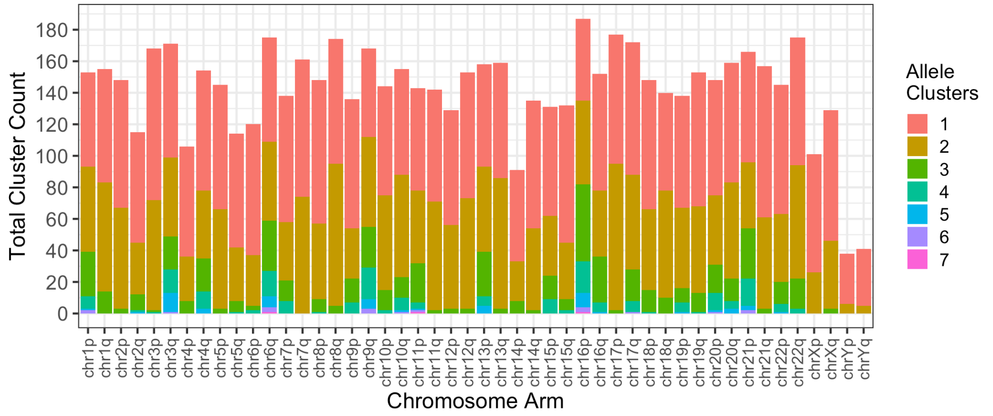


**C**

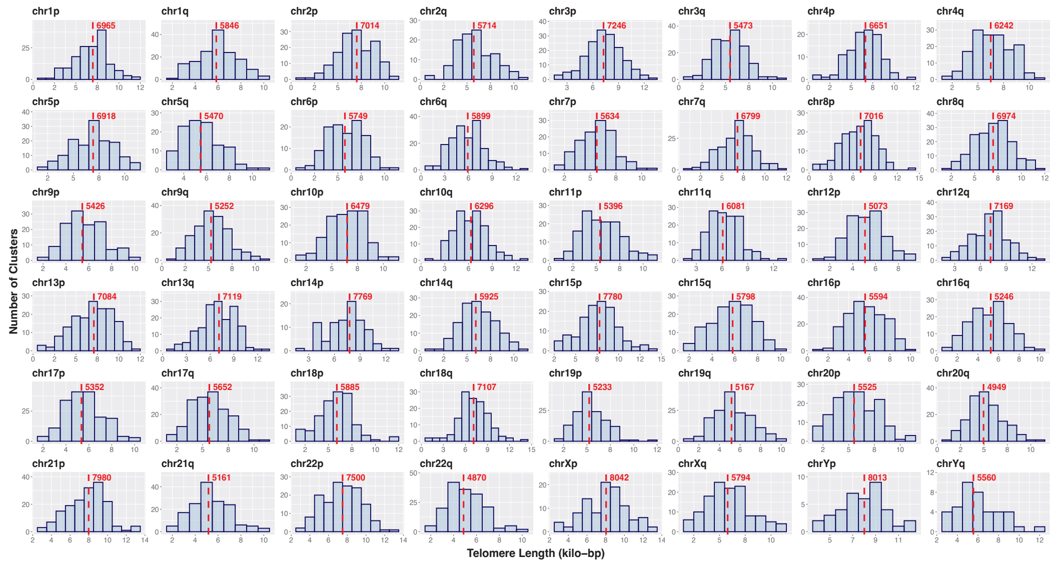

**D**

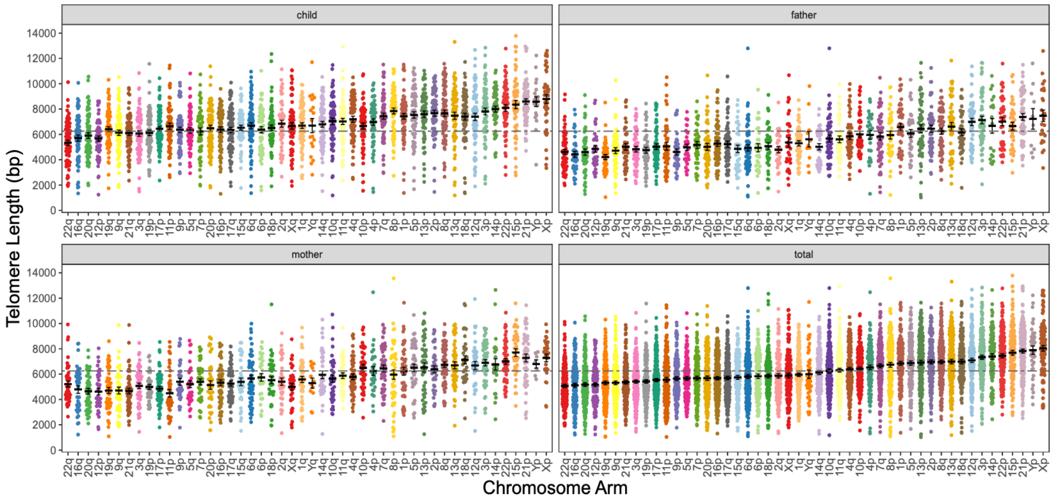


**E**

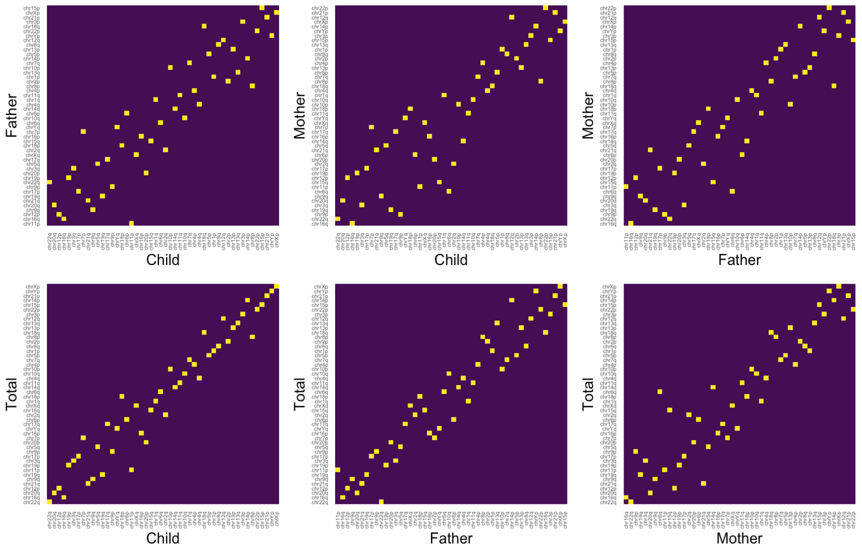


**Supplementary Figure 1: Chromosome arm-specific TL distributions and rank-order correlations in GA4K trios.**

(A) Telomere candidate read counts at each arm chromosome arm for all participants in GA4K trios.

(B) The number of reported telomeric allele clusters per chromosome arm across all individuals in GA4K trios.

(C) Chromosome arm-specific TL in GA4K trio cohort (n = 225). Histograms showing TLs distributions, with red vertical lines indicating peak frequency in base pairs (bps).

(D) Chromosome arm-specific TL distributions across family members. TL distributions per chromosome arm in children, fathers, mothers, and combined cohort (GA4K trios, n = 225), shown in the arm order ranked by median TL across the entire cohort. This visualization allows assessment of whether TL rank order is preserved across family roles.

(E) Rank-order comparison heatmaps showing pairwise correlations between median TLs across 46 chromosome arms for child–father, child–mother, father–mother, and total comparisons. High correlation across roles confirms that arm-specific TL ranks are broadly conserved within families.

**A**

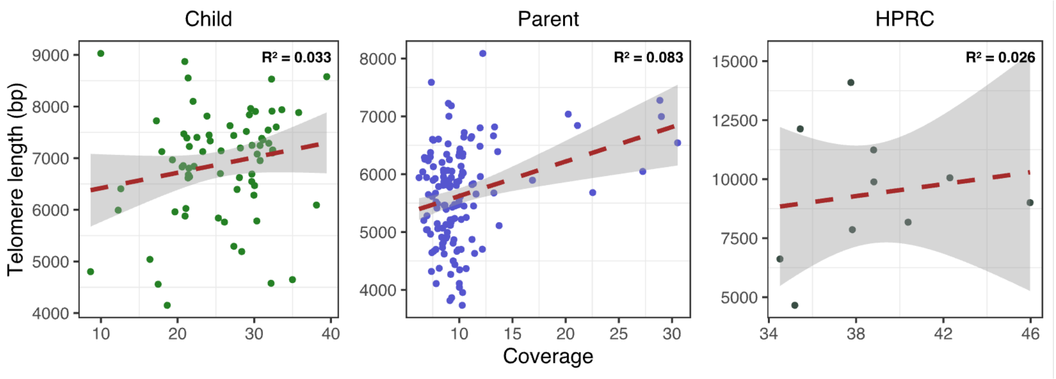


**B**

```
Coefficients:
                 Estimate Std. Error t value Pr(>|t|)
(Intercept)      288.2737   965.3433   0.299  0.76624
meanTL_mother      0.5023     0.1165   4.313 5.99e-05 ***
meanTL_father      0.5018     0.1063   4.718 1.43e-05 ***
age_child        -45.4357    14.8788  -3.054  0.00335 **
age_father_birth -22.0141    27.3958  -0.804  0.42477
age_mother_birth  59.0650    32.2409   1.832  0.07184 .
coverage_child     4.9903    13.1909   0.378  0.70651
---
Signif. codes:  0 ‘***’ 0.001 ‘**’ 0.01 ‘*’ 0.05 ‘.’ 0.1 ‘ ’ 1

Residual standard error: 663.8 on 61 degrees of freedom
Multiple R-squared:  0.5956,    Adjusted R-squared:  0.5558
F-statistic: 14.97 on 6 and 61 DF,  p-value: 1.928e-10
```

**Supplementary Figure 2: Impact of HiFi-GS sequencing coverage on telomere length prediction.**

(A) Examination of sequencing coverage impact across different sample groups. The relationship between TL and HiFi-GS sequencing coverage is shown for GA4K children and parents (68 trios, n = 204), and a comparative HPRC sample group (n = 10). The trend line is displayed for each sample subgroup, with the gray shading representing the standard error of the estimate.

(B) Summary of multiple linear regression analysis testing the inclusion of child sequencing coverage as a predictor in the model, showing that sequencing coverage is not a significant predictor of children’s MTL.

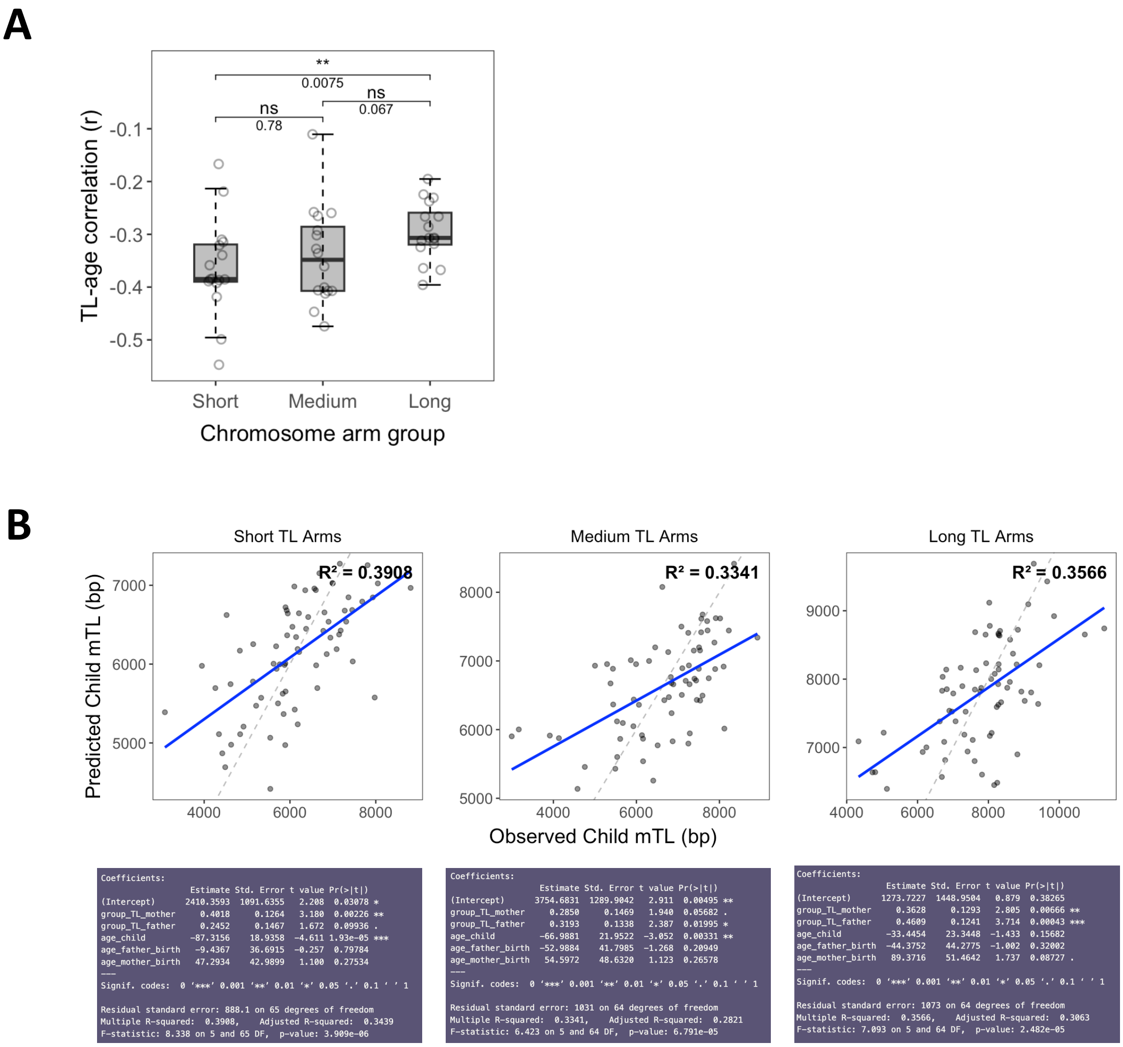


**Supplementary Figure 3: Impact of chromosome arm grouping on child's mean telomere length prediction.**

(A) Correlation between telomere length and age varies across chromosome arm groups. Each data point represents the Pearson correlation coefficient (r) between TL and age for a given chromosome arm, grouped by telomere length category (Short, Medium, Long). Boxplots show the distribution of correlation values within each group. Statistical comparisons between groups were performed using the Wilcoxon rank-sum test; adjusted p-values are indicated.

(B) Actual versus predicted child TL by chromosome arm group (short TLs, medium TLs, long TLs). Regression performance summaries are shown below.

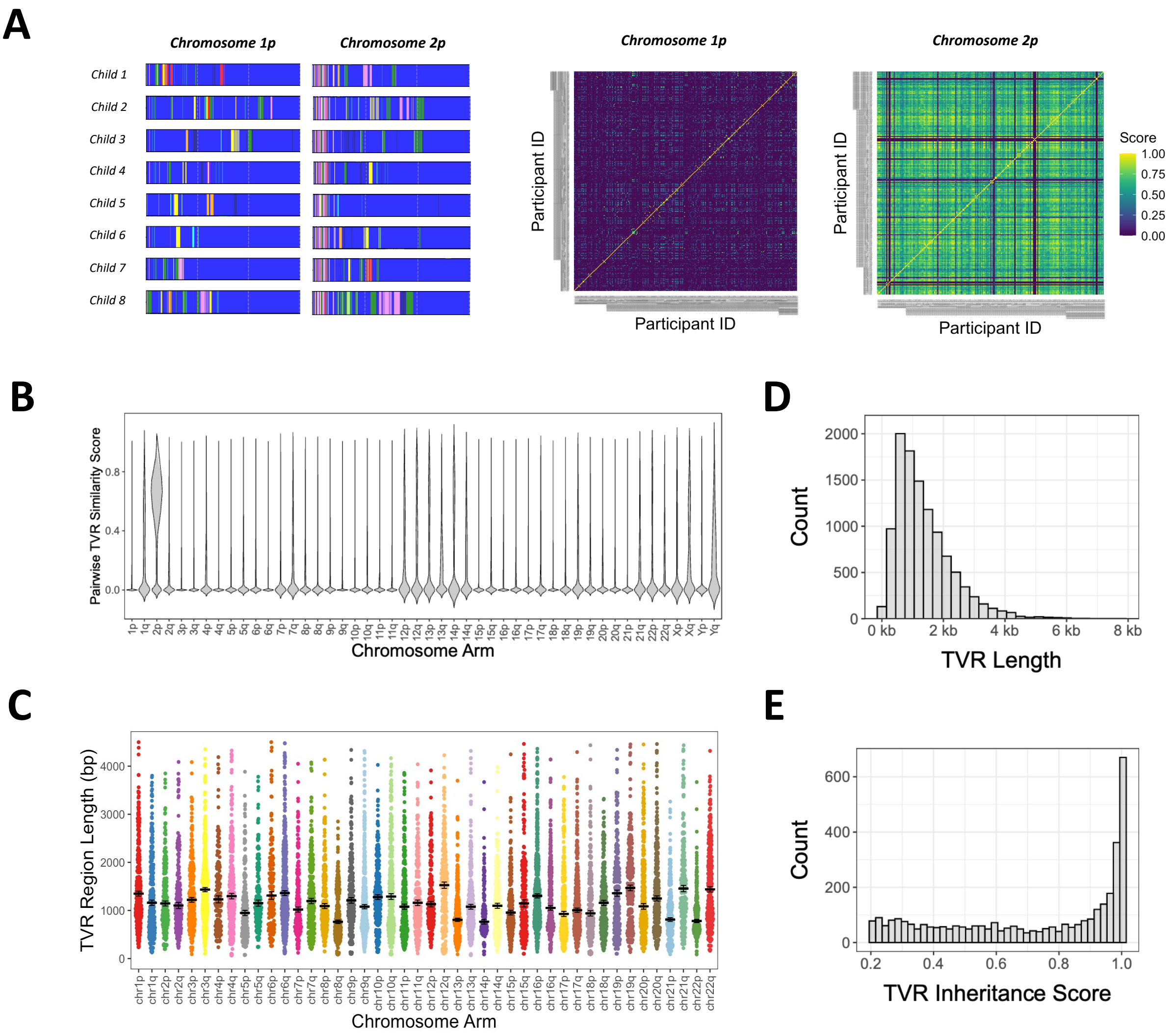


**Supplementary Figure 4: TVR characterization across chromosome arms and inheritance scores in GA4K trios.**

(A) (Left) TVR consensus plots from eight unrelated individuals show visually similar repeat organization at 2p, whereas 1p displays more variable patterns. (Right) Pairwise similarity matrices comparing all reported TVR alleles for each arm across the GA4K cohort show higher similarity scores at 2p (n = 292) than at 1p (n = 281), suggesting greater structural conservation at 2p across individuals.

(B) Violin plots showing the distribution of pairwise TVR similarity scores across all chromosome arms, excluding self-comparisons and within-family comparisons.

(C) TVR region length distributions by chromosome arm.

(D) Distributions of TVR lengths across all telomeric alleles.

(E) The distribution of inheritance scores, showing the pairwise similarity between child and parent TVR consensus alleles across all parent-child pairs. A similarity score of ≥ 0.7 was used as the cutoff for calling an allele inherited.

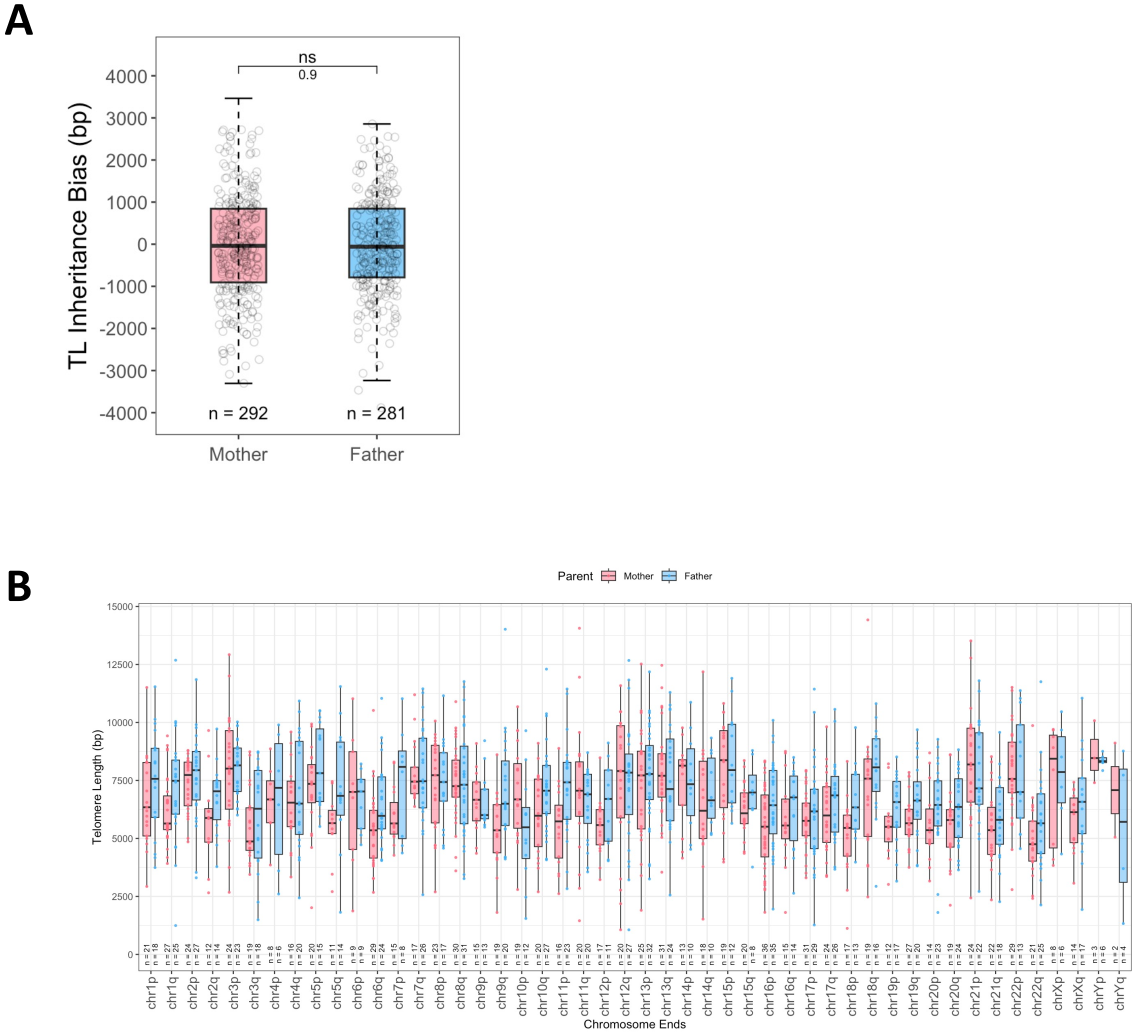


**Supplementary Figure 5: Telomeric allele inheritance patterns stratified by parental origin.**

(A) Telomere drive analysis separated by parental role. Inheritance bias was calculated as the difference between the transmitted allele TL and the average TL of both alleles from the transmitting parent at the same chromosome arm. No consistent bias was observed in either maternal or paternal transmissions.

(B) Comparison of inherited telomeric alleles measured in children across all trios, shown separately for paternally and maternally transmitted alleles at each chromosome arm.

### Supplementary Table

| Acrocen Chr | Sex Chr | Rm rlen <1000 | (Intercept) | **mTL_mother** | **mTL_father** | **age_child** | **age_father_ childbirth** | **age_mother _childbirth** | **coverage_ child** | **Multiple R-squared** | **Adjusted R-squared** | **Model p-value** | |
|---|---|---|---|---|---|---|---|---|---|---|---|---|---|
| | | T | 0.76624 | 5.99e-05 *** | 1.43e-05 *** | 0.00335 ** | 0.42477 | 0.07184 . | 0.70651 | 0.5956 | 0.5558 | 1.93E-10 | |
| | | F | 0.33206 | 0.00019 *** | 0.00045 *** | 0.00521 ** | 0.12692 | 0.04091 * | 0.71402 | 0.5315 | 0.4854 | 1.39E-08 | |
| | | T | 0.72246 | 2.98e-05 *** | 9.97e-06 *** | 0.00330 ** | 0.41138 | 0.06736 . | | 0.5947 | 0.562 | 4.54E-11 | **Used model** |
| | | F | 0.28829 | 6.85e-05 *** | 0.00041 *** | 0.00522 ** | 0.11535 | 0.03653 * | | 0.5304 | 0.4926 | 3.68E-09 | |
| | exclu | T | 0.67851 | 1.45e-05 *** | 2.30e-05 *** | 0.00256 ** | 0.36082 | 0.06617 . | | 0.5915 | 0.5585 | 5.75E-11 | |
| | exclu | F | 0.24037 | 3.68e-05 *** | 0.00079 *** | 0.00403 ** | 0.08685 . | 0.03286 * | | 0.5275 | 0.4894 | 4.43E-09 | |
| exclu | exclu | T | 0.24165 | 3.18e-06 *** | 0.00081 *** | 0.00074 *** | 0.34877 | 0.14088 | | 0.5681 | 0.5333 | 3.03E-10 | |
| exclu | exclu | F | 0.04736 * | 2.50e-05 *** | 0.01520 * | 0.00209 ** | 0.07331 . | 0.06530 . | | 0.4851 | 0.4436 | 5.65E-08 | |

Significance codes: 0 '***' 0.001 '**' 0.01 '*' 0.05 '.' 0.1 ' ' 1.

**Supplementary Table 1: Impact of varied TL calculation methods and predictors on multiple linear regression model performance.**

Summary of regression model optimization. We tested different telomere length (TL) calculation strategies and predictor combinations to evaluate multivariate model performance. Averaging strategies compared TL calculated over all chromosome arms, autosome-only arms, and arms excluding acrocentric chromosomes. We also tested the effect of filtering outlier candidate reads with TL <1000 bp. Additionally, we evaluated the influence of child HiFi-GS sequencing coverage by including it as a sixth predictor in the regression model (MTL_child ~ MTL_mother + MTL_father + Age_child + Age_mother_conception + Age_father_conception + Coverage). Each model performance was assessed based on adjusted $R^2$ and model p-values. The final model was selected based on the overall statistical significance and best predictive fit.

**Supplementary Table 2: Summary of clinical findings, genetic variants, and HiFi-GS TL measurements in three probands.**

| **Proband Label** | **Age at Sample Collection** | **Sex** | **Flow-FISH TL (lymphocyte)** | **Long-read MTL** | **Diagnosis** | **Key variant(s)** |
|---|---|---|---|---|---|---|
| Proband_A | 1 y | M | 5.4k | 4433 | DC (dual) | TERT G1063S + chr14 del |
| Proband_B | 2 y | F | 8.1k | 6500 | X-linked DC | DKC1 G326V |
| Proband_C | 2 mo | F | 10.6k | 8678 | NDD + cytopenia | DOT1L E123K (VUS) |

| | Role | # arms with TL calls |
|---|---|---|
| At least 1 allele has TL call | Child | 3026 |
| | Mother | 1909 |
| | Father | 1961 |
| At least 2 alleles have TL calls | Child | 1553 |
| | Mother | 563 |
| | Father | 567 |

**Supplementary Table 3:** Summary of the number of chromosome arms with telomeric allele calls across trios. For each family member (child, mother, and father), we report how many chromosome arms had at least one telomeric allele call. Arms with two or more called alleles are also indicated.

| Parent-child TVR inheritance tracing | | # shared arms with TL calls | Correct assignments | Incorrect assignments | Success rate (%) | Error rate (%) |
|---|---|---|---|---|---|---|
| Mother-child | 1 allele | 1593 | 849 | 47 | 53.30% | 2.95% |
| | 2 alleles | 437 | 313 | | 71.62% | |
| Father-child | 1 allele | 1618 | 807 | 48 | 49.88% | 2.97% |
| | 2 alleles | 430 | 299 | | 69.53% | |

**Supplementary Table 4: Summary of TVR-based inheritance tracing for each parent-child pair.**

Performance summary of TVR-based allele tracing in trios. Shared arms are defined as chromosome arms where both the parent and the child have at least one allele with a TL call. We report the number of correct inheritance assignments, incorrect arm assignments, success rate, and estimated error rate. The estimated error rate is defined as the proportion of cases where the top-scoring TVR match comes from a different chromosome arm than expected. Similar to Table S2, “2 alleles” refers to shared arms where both the parent and child have two telomeric allele calls.
